# Temporal stability and genetic diversity of 48-year old T-series phages

**DOI:** 10.1101/2020.09.24.312751

**Authors:** Dinesh Subedi, Jeremy J. Barr

## Abstract

T-series phages have been model organisms for molecular biology since the 1940s. Given that these phages have been stocked, distributed, and propagated for decades across the globe, there exists the potential for genetic drift to accumulate between stocks over time. Here we compared the temporal stability and genetic relatedness of laboratory-maintained phage stocks with a T-series collection from 1972. Only the T-even phages produced viable virions. We obtained complete genomes of these T-even phages, along with two contemporary T4 stocks. Performing comparative genomics, we found 12 and 16 nucleotide variations, respectively, in the genomes of T2 and T6; whereas there were ~172 nucleotide variations between T4-sublines when compared with NCBI RefSeq genome. To account for the possibility of artefacts in the NCBI RefSeq, we used the 1972 T4 stock as a reference and compared genetic and phenotypic variations between T4-sublines. Genomic analysis predicted nucleotide variations in genes associated with DNA metabolism and structural proteins. We did not however, observe any differences in growth characteristics or host range between the T4-sublines. Our study highlights the potential for genetic drift between individually maintained T-series phage stocks, yet after 48-years, this has not resulted in phenotypic alternations in these important model organisms.

**Importance:** T-series bacteriophages have been used throughout the world for various molecular biology researches, which were critical for establishing the fundamentals of molecular biology – from the structure of DNA to advanced gene-editing tools. These model bacteriophages help keep research data consistent and comparable between laboratories. However, we observed genetic variability when compared contemporary sublines of T4-phages to a 48-year old stock of T4. This may have effects in the comparability of results obtained using T4 phage. Here, we highlighted the genomic differences between T4 sublines and examined phenotypic differences in phage replication parameters. We observed limited genomic changes but no phenotypic variations between T4 sublines. Our research highlights the possibility of genetic drift in model bacteriophages.

## Introduction

Bacteriophages, viruses that infect bacteria, were discovered in the early 20^th^ century. The antimicrobial properties of phages initially sparked the interest of those early phage-pioneers and they were quickly used to treat bacterial infections, such as dysentery and cholera (Rohwer and Segall 2015). However, the ambiguous efficacy of phage therapy, along with the introduction of more efficient chemotherapeutics, led to the subsequent decline in interest in their use as therapeutic agents (Salmond and Fineran 2015). Nevertheless, since their discovery, phages have become key model organisms to understand various aspects of modern molecular biology. For example, understanding of the basis of mutation (Luria and Delbruck 1943), recombination (Luria and Human 1952), the genetic nature of DNA and its replication (Crick et al 1961), and the sequencing of genes and genomes (Sanger et al 1977) were all founded upon phage biology. Furthermore, the study of phage prokaryote resistance mechanisms led to the discovery of the CRISPR/Cas system, which has become a key technique for targeted mutagenesis and gene editing (Barrangou et al 2007, Doudna and Charpentier 2014). Nowadays, amidst the looming threat of antibiotic resistance, phage therapy has regained global attention (Gordillo Altamirano and Barr 2019).

One of the most important historical advancements in phage biology was the introduction of the T-series phages in the 1940s by Delbruck and colleagues’– the so-called ‘phage group’. This enabled phage researchers to make comparison of results between different laboratories, which was previously unsystematic, due to the data coming from a random collection of phage-host combinations. T-series phages constitute a collection of seven virulent phages (T1-T7; T=type), which were described based on their ability to lyse *Escherichia coli* B (Demerec and Fano 1945, Luria and Delbruck 1943). Although this classification was merely based on the lytic activity of the group, T2, T4, and T6 (T-even series) turned out to be similar; morphologically, antigenically, and genetically (Abedon 2000). T-even phages are classified as members of the *Myoviridae* family with their characteristic contractile tail. The genomes of T-even phages contain between 160-170 kbp dsDNA, which has 5-hydroxymethyl-cytosine in place of cytosine (Kutter and Wiberg 1968). In contrast, T-odd phages (T1, T3, T5 and T7) are highly variable between each other. They have a relatively simple non-contractile tail and, unlike the T-even series, contain dsDNA with the usual four nucleotides. Genome size is also diverse amongst the T odd phages (T1=48 Kbp, T3=38 Kbp, T5=121 Kbp, T7=40 Kbp). The traditional T-series phages are still in use today worldwide. They have been distributed, maintained, and propagated across many different laboratories and repositories for decades. However, questions remain regarding the stability and genetic drift of model T-series bacteriophages over time and how these changes have shaped bacterial hosts past and present.

We, therefore, sought to examine the temporal stability of T-series phages in prolonged storage and compare the genetic relatedness of different laboratory-maintained T4 phage stocks. We obtained a near-complete T-series phage stock (T1, T2, T3, T4, T6 and T7 – notably T5 was missing) that had been prepared by Robert E. Hancock and co-workers at the University of Adelaide, Australia in 1972 (Hancock and Reeves 1975, Hancock et al 1976). The T-series lysates had been purified and stored in chloroform-sealed glass ampules (Fig 1). To evaluate the stability of these phages, we quantified active phages in the ampules by standard top-agar assays. To compare the genetic divergence of the phages, we obtained two contemporary stocks of T4 (hereafter referred to as T4 sublines for simplicity) from phage laboratories in Australia and the USA. We then compared the genetic and phenotypic differences of these sublines to Hancock’s 1972 stock of T4.

**Figure 1:**
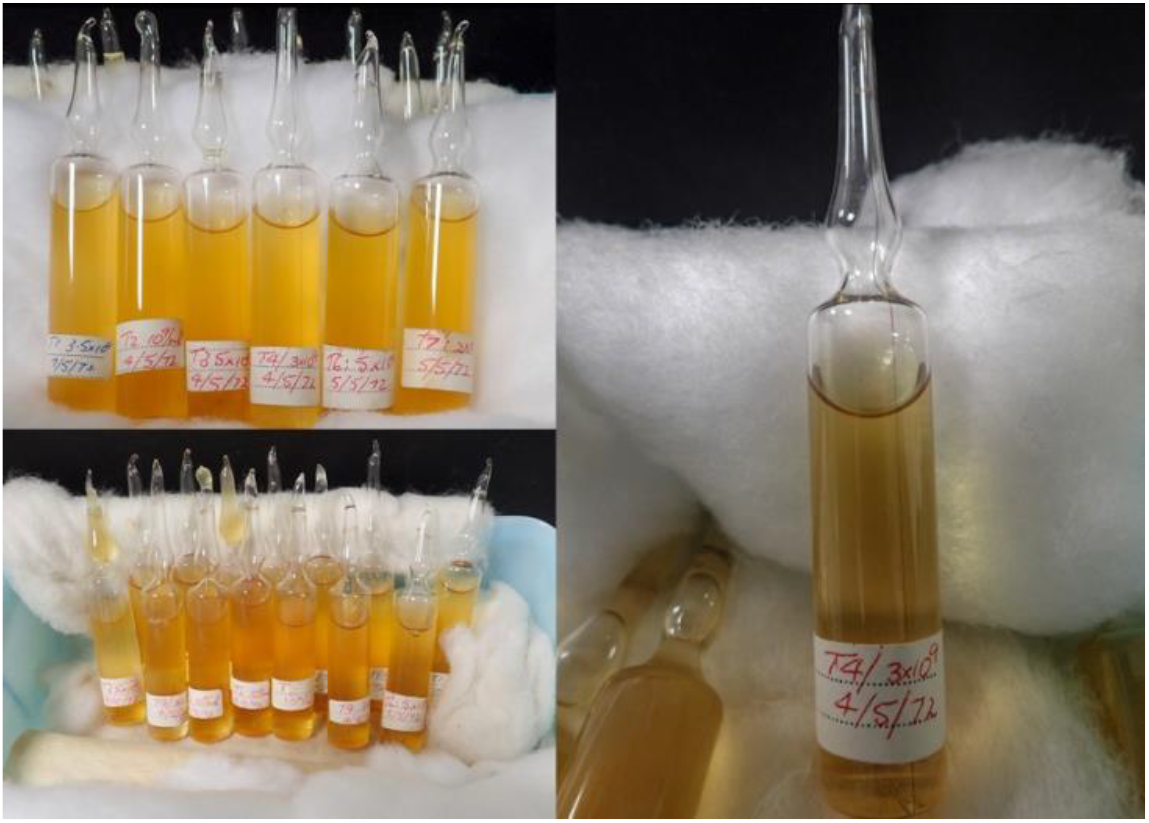
Photographs of historical stocks of T-series phages, stored in 1972. Each glass ampule contained 10 mL lysate stored in chloroform-sealed glass ampules at a titre of 1-5 × 10^9^ plaques forming unit per mL. The lysates have been stored at 4 °C since 1972.

## Methods

### Bacteriophage stock

The stock of T-series bacteriophages in this study was obtained from Prof. Peter Reeves, from The University of Sydney, Australia. The stock comprised lysates of T1, T2, T3, T4, T6 and T7 phages, purified and stored in sealed glass ampules with chloroform in 1972. These phages were used in several studies by Robert E. Hancock in the 1970s (named T#-Hancock henceforth) (Hancock and Reeves 1975, Hancock et al 1976, Hancock and Reeves 1976, Hancock and Braun 1976). T1 phage was not used in this study due to its potential contamination hazard for our laboratory. These stocks have been stored at cold room (~4 °C) since the 1970s. In addition, we used two contemporary strains of T4; received from The Bacteriophage T4 lab at the Catholic University of America led by Prof. Venigalla B. Rao (named T4-Rao henceforth) and the phage collection of our lab (named T4-Barr henceforth).

### Bacteriophage revival and propagation

Lysates from the old stocks were each recovered in 15 mL falcon tubes. 1 mL of lysate was mixed with susceptible hosts (*E. coli* B for T2/3/4/6 and *E. coli* BL21 for T7) and plated onto Lysogeny Broth (LB) agar plates using the soft agar overlay method, followed by incubation for 18-24 h at 37 °C. Each plating was done in duplicate and repeated with lysates from at least three different vials. In instances of no plaque formation, amplification was attempted using the original lysate and its host in broth culture, then replated. Lysates with viable phages were propagated following the Phage-on-Tap protocol (Bonilla et al 2016).

### Phage DNA extraction and sequencing

High titre lysate (>10^9^PFU/mL) was used for DNA extraction. To remove host’s nucleic acid contamination, lysates was treated with DNase (1 mg/ml) and RNase A (12.5 mg/ml) for 2 h at 37 °C, followed by an inactivation step of 5 min at 75 °C. DNA was extracted using Norgen phage DNA extraction kit (Norgen Biotek, Ontario, Canada), following the manufacturer’s instructions. The extracted DNA was vacuum dried into a pellet for transport. Sequencing was performed using the Illumina® HiSeq 150 bp paired-end platform at the Genewiz® facilities (Suzhou, China).

### Bioinformatics analysis

Illumina Hiseq platform generated five to six million raw reads from each sample. Trimmomatic v0.39 (Bolger et al 2014) was used to remove adaptor sequences and to truncate reads of quality less than 15 in a sliding window of 4 bp. Raw reads were assembled using Unicycler v0.43 (Wick et al 2017) with a flag of minimum contig length of 1000 bp, which produced a single and complete genome. We aligned NCBI RefSeqs (T2 nucleotide accession; LC348380; T4 nucleotide accession: AF158101; T6 nucleotide accession: MH550421) with the assembled genomes using Mauve genome alignment (Darling et al) and the results from the alignment were used to re-arrange nucleotides to match with the start position of NCBI RefSeqs. The genomes were then annotated manually copying the annotation from their respective NCBI RefSeqs using Geneious v 9.1.8 (Kearse et al), provided a sequence similarity of coding region of at least 98% followed by manual curation where required.

To identify nucleotide variants, the filtered raw reads were mapped to either NCBI RefSeqs or our assembled genomes using Snippy v4.2 (https://github.com/tseemann/snippy) with the setting of the minimum number of reads covering a site to be considered at 100 and the minimum VCF variant call quality also at 100. To compare nucleotide identity, the complete genomes were aligned by pairwise sequence alignment as implemented in Pyani (https://huttonics.github.io/pyani/). Average nucleotide identity (ANI) percentage obtained from the analysis was converted into matrix and visualised using pheatmap (Kolde 2012) in R package.

#### Nucleotide accessions

The nucleotide sequence of T4-Hancock is available in NCBI GenBank accession number MT984581. Raw sequence data and alignment files are available in NCBI BioProject ID PRJNA662192.

### Determination of T4 growth characteristics

The growth characteristics of all three T4 phages were examined using hosts *E. coli* B and *E. coli* K12. Firstly, relative efficiency of platting between hosts was examined. Each lysate was serially diluted to a titre of ~100 PFU/mL, and 500 μL was used in a soft agar overlay with each host. After overnight growth, PFUs were counted and recorded and the relative efficiency of plating on *E. coli* K12 was calculated as the ratio of PFU in *E. coli* K12 to PFU in *E. coli* B. Each experiment was performed in duplicate and repeated three times.

Next, one-step growth curve experiments were performed on *E. coli* B and *E. coli* K12 separately with each T4 subline. Bacteria from overnight broth cultures were diluted 1:20 in LB and allowed to grow until an optical density 600 nm (OD600) reached 0.2 (~2 h), which was then infected with phage at a multiplicity of infection (MOI) of 0.01. Phage was allowed to adsorb for 5 min at 37 °C with orbital agitation at 120 rpm. The mixture was then pelleted (4,000 g, 2 min, room temperature), resuspended in fresh LB broth and taken back to incubation at 37 °C and 120 rpm. Samples (100 μL) were repeatedly taken every 5 or 10 min for a total period of 1 h, transferred into chloroform-saturated PBS, serially diluted, and plated to determine PFU. Number of bacteria in initial inoculum was determined by plate count. PFU per infected cells was calculated by dividing PFU by initial density of bacteria. The experiment was repeated on at least three different occasions.

Bacterial lysis by phage was determined from sequential OD readings obtained using microtiter plate reader. Bacterial cultures at exponential growth were obtained as described above. Each culture was added to individual wells of a 96-well microtiter plate and mixed with phage lysate at 0.01 MOI. The plate was incubated at 37 °C in a microplate reader (Epoch™ Microplate Spectrophotometer, BioTek, Winooski, VT, USA) with continuous shaking and the data was recorded every 5 min.

To gauge the host range of each T4-subline, we used 11 different clinical and laboratory isolates of *E. coli* and the standard spot plate assay (Kutter 2009). 10 μL of phage lysate was spotted on the surface of agar plate seeded with host bacteria. The zone of clearance on the spotted area was recorded as a positive lytic activity.

### Graphing, data presentation and statistical analysis

All data were analysed and visualised using Graphpad Prism v8. Average and standard deviation of all the replicates were calculated and compared using t-test to obtain p values. To infer statistical significance, the threshold was set at values of p < 0.05.

## Results

### Stability of T-series phages in prolonged storage

To examine the viability of the 48-year old T-series phage stocks, we plated 1 mL of lysate from T2, T3, T4, T6 and T7 phages with their respective hosts using the soft-agar overlay technique. We did not open or plate T1 phage vials due to concerns with its persistence and history of contaminating laboratory stocks of *E. coli*. The titres in plaque-forming units (PFU) per mL were recorded from at least three different vials for each strain (Table 1). We did not observe any plaques from T3 and T7 lysates. To verify whether there were any active phage particles in the T3 or T7 stocks remaining at low titre, we propagated entire vial of the original lysates with *E. coli* B and *E. coli* BL21, respectively, overnight in an attempt to recover viable phages. However, no phages could be recovered following overnight amplification. Lysates of T2 and T4 showed between 4 to 6 plaques per mL on top-agar, while T6 phage had between 10^4^ to 10^5^ PFU/mL (Table 1). Our results revealed that T-even series could be revived from the lysates stored approximately 48-years ago, while T-odd (T3 and T7) series were not able to be recovered from the 1970’s stock.

**Table 1:** Growth of T-series phages from 48-year old stocks (PFU = plaque forming unit)

| Bacteriophage | Ampule 1 | Ampule 2 | Ampule 3 | Average PFU/mL |
| --- | --- | --- | --- | --- |
| | PFU/mL | PFU/mL | PFU/mL | (mean ± SD) |
| T2 | 4 | 3 | 3 | $3.34 \pm 0.47$ |
| T3 | No growth | No growth | No growth |  |
| T4 | 3 | 6 | 2 | $3.67 \pm 1.7$ |
| T6 | $4.2 \times 10^4$ | $3.1 \times 10^5$ | $6.2 \times 10^4$ | $1.38 \times 10^5 \pm 1.21 \times 10^5$ |
| T7 | No Growth | No Growth | No Growth |  |

### Genomic characteristics of T-even series

The complete genomes of 1972 stocks of T2, T4 and T6 phage, henceforth referred to as ‘Hancock’ phages, were obtained using Illumina HiSeq. Raw reads of each genome were aligned with the corresponding NCBI RefSeq to examine single nucleotide polymorphisms (SNPs) and insertion deletion variations (indels) using Snippy v4.2.0. We observed 12 and 16 variants (SNPs + indels) in the T2-Hancock and T6-Hancock phages, respectively, when compared with NCBI Refseq (T2 nucleotide accession; LC348380; T6 nucleotide accessions: MH550421) with the nucleotide variants being distributed throughout the genome (Table 2, supplementary file). Comparatively, when we investigated the T4-Hancock phages we found 172 nucleotide variants compared to the NCBI T4 Refseq (Nucleotide accession AF158101). The reason for this high nucleotide variation was not immediately clear. The original T4 genome sequence in the NCBI database was last updated in 2003 and had been completed using traditional sequencing methods involving cloning and PCR (Miller et al 2003). To examine whether this difference arose due to artefacts in T4’s RefSeq, we compared genome sequences of two contemporary T4 sublines from The Barr Lab, Monash University, School of Biological Sciences, Australia (henceforth referred to as T4-Barr) and from The Bacteriophage T4 Lab, led by Venigalla B. Rao, The Catholic University of America, Washington D.C., USA (henceforth referred to as T4-Rao) to the NCBI Refseq. We observed that T4-Barr and T4-Rao had similar nucleotide variations of 172 and 175 respectively, compared with the T4 RefSeq (Table 2, supplementary file). This led us to conclude that there may be artefacts in the NCBI RefSeq genome, and we therefore used T4-Hancock as a historical reference genome in our subsequent analysis.

**Table 2:** Genome size and nucleotide variants of T-series phages compared with NCBI RefSeq genomes.

| Strains | Genome<br>Size (bp) | NCBI RefSeq<br>Accession no. (genome size) | Nucleotide variants<br>Total (SNPs +indels) |
| --- | --- | --- | --- |
| T2-Hancock | 163,826 | LC348380 (163,825bp) | 12 (8+4) |
| T4-Hancock | 168,908 | AF158101 (168,903bp) | 172 (97+75) |
| T4-Barr | 168,921 | AF158101 (168,903bp) | 172 (99+73) |
| T4-Rao | 168,922 | AF158101 (168,903bp) | 175 (100+75) |
| T6-Hancock | 168,702 | MH550421(168,706bp) | 16 (8+8) |

### Comparative genomics of T4 sublines

To compare the complete genomes of T-even phages, we assembled the Illumina HiSeq reads using Unicycler v0.4.3 with the setting of minimum contig length 1000bp, which produced a single contig of a complete genome for each strain. The genome size of T-even series of Hancock strains was broadly similar to the respective NCBI Refseq (Table 2). The complete genomes were aligned by pairwise sequence alignment as implemented in Pyani (https://huttonics.github.io/pyani/) to obtain the average nucleotide identity (ANI). Analysis revealed that genomes of T-even phages were closely related with each other, with more than 96% nucleotide sequence similarity between T2, T4 and T6 phages (Supplementary Figure 1). When compared between T4 sublines, we observed that T4-Barr and T4-Rao showed higher percentage similarity than when compared with T4-Hancock (Figure 2).

**Figure 2:**
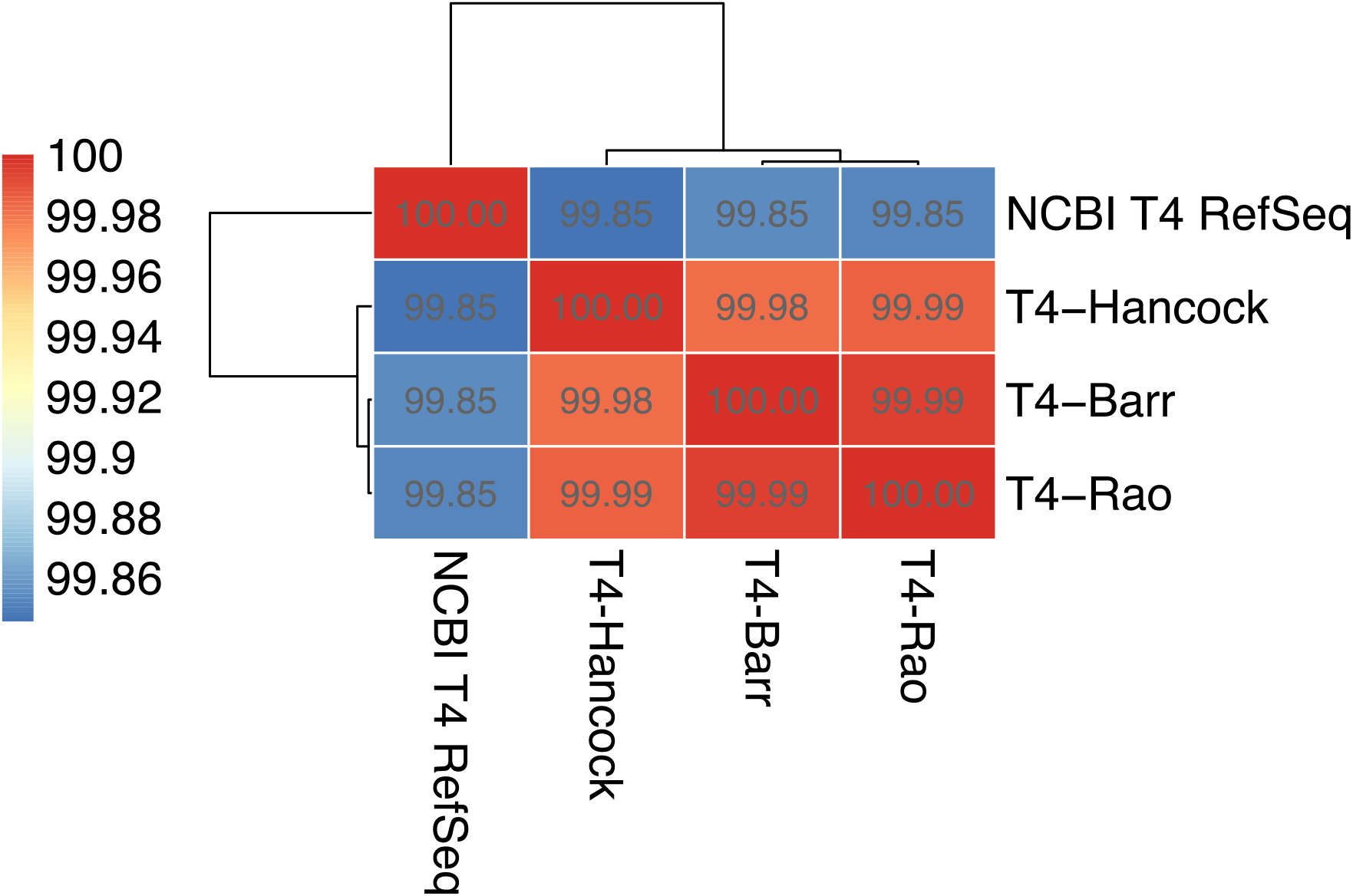
Average nucleotide identity matrix of T4 sublines. Dendrograms represent sequence similarity between T4 genomes. The bar on the left represents colour code for percentage similarity (NCBI T4 RefSeq refers to the GenBank accession number AF158101).

Next, we assessed the nucleotide variations (SNPs and indels) between T4 sublines. To obtain an unbiased annotation, we manually annotated the complete genome of each subline using information from NCBI T4 Refseq using an identity threshold of at least 98%. T4-Barr had two insertions (frameshift) and six SNPs (five missense and one synonymous), when compared to T4-Hancock. Our analysis showed five additional variants (one insertion and four missense SNPs) in T4-Rao, taking the total number of variants to 13 (Table 3).

**Table 3:** Nucleotide variants in T4-Barr and T4-Rao compared to T4-Hancock as the reference. Bold typeface denotes variants present in T4-Rao but absent in T4-Barr.

| Gene | product | Variants |  | Type of variants | Function categories** |
| --- | --- | --- | --- | --- | --- |
|  |  | Nucleotides | Amino acids |  |  |
| rIIA | protector from prophage-induced early lysis | 222A>C | Glu74Asp | missense | Auxiliary |
| 39 | topoisomerase II, large subunit, N-terminal region | 475A>G | Ile159Val | missense | Essential |
| dda | DNA helicase | 1174G>A | Ala392Thr | missense | Auxiliary |
| 55.2 | hypothetical protein | 123A>G | Gln41Gln | synonymous |  |
| rI.-1 | **MobD.6 hypothetical protein** | **179dupG** | **Glu61fs** | **frameshift** |  |
| tk.4 | conserved hypothetical protein | 106A>G | Ser36Gly | missense |  |
| 7 | baseplate wedge initiator | 776T>C | Phe259Ser | missense | Essential |
|  | *Intragenic region |  |  |  |  |
| 18 | **tail sheath protein** | **326G>A** | **Ser109Asn** | **missense** | **Essential** |
| alt | **Alt RNA polymerase ADP-ribosylase** | **691C>T** | **Arg231Cys** | **missense** | Auxiliary |
| 34 | long tail fiber, proximal subunit | 1151C>T | Pro384Leu | missense | Essential |
| 37 | **long tail fiber, distal subunit** | **1250G>A** | **Gly417Asp** | **missense** | **Essential** |
| arn | Arn inhibitor of MrcBC restriction endonuclease (anti-restriction nuclease) | 258dupA | Leu87fs | frameshift | Auxiliary |
| 52 | **topoisomerase II, medium subunit** | **757G>A** | **Asp253Asn** | **missense** | **Essential** |

\* an insertion of GAAGCTTTGGCCC at the intergenic region at 93,397bp \*\*assigned according to Miller, 2003 (Miller et al 2003)

Frameshift mutations were observed in an auxiliary gene (*arn*) and a hypothetical protein (MobD.6). *Arn* encodes an anti-restriction endonuclease that inhibits *E. coli* host’s endonuclease activity (Dharmalingam et al 1982). Our manual annotation of T4-Hancock showed that Arn consists of 94 amino acids, which is slightly different from that of NCBI T4 Refseq (92 amino acids). Interestingly, duplication of nucleotide A at position 258 of *arn* resulted in a stop codon at position 276, truncating the protein length to 92 amino acids in T4-Barr and T4-Rao compared to 94 amino acids in T4-hancock. The frameshift mutation in MobD.6 also caused gained of a premature stop codon at position 180 (of 387 bp), potentially resulting in a loss-of-function mutation in this hypothetical gene.

On the other hand, many missense variations were observed in genes associated with essential structures such as two units of topoisomerase II (gp39 and gp52), which helps in DNA metabolism, and the baseplate wedge initiator (gp7), tail sheath protein (gp18) and long tail fibres (gp34 and gp37), all associated with phage adsorption to the bacterial host. Out of the five additional variants in T4-Rao, three were observed in essential genes; tail sheath protein (gp18), distal subunit of long tail fibre (gp37) and medium subunit of topoisomerase (gp52).

### Growth characteristics of T4 sublines in *E. coli* B and *E coli* K12

We then sought to assess whether the identified mutations had effects on phage replication variables. For these experiments, we used two hosts: *E. coli* B and the closely related strain *E. coli* K12. T4 adsorbs to these two *E. coli* strains using different receptors. In *E. coli* K12, T4 uses lipopolysaccharide (LPS) and outer membrane protein C (OmpC) as receptors. However, in *E coli* B, which harbours a deletion in *ompC*, T4 uses LPS as the sole receptor (Washizaki, Yonesaki, & Otsuka, 2016; Yu & Mizushima, 1982). Firstly, we performed an efficiency of plating (EOP) assay to examine relative titres of T4 sublines on *E. coli* K12 compared to the original host of propagation, *E. coli* B. Our results showed that the relative EOPs were not significantly different between any of the three sublines of T4. However, T4-Rao had slightly lower EOPs (1.012) compared to T4-Hancock (1.053) and T4-Barr (1.068) (Figure 3).

**Figure 3:**
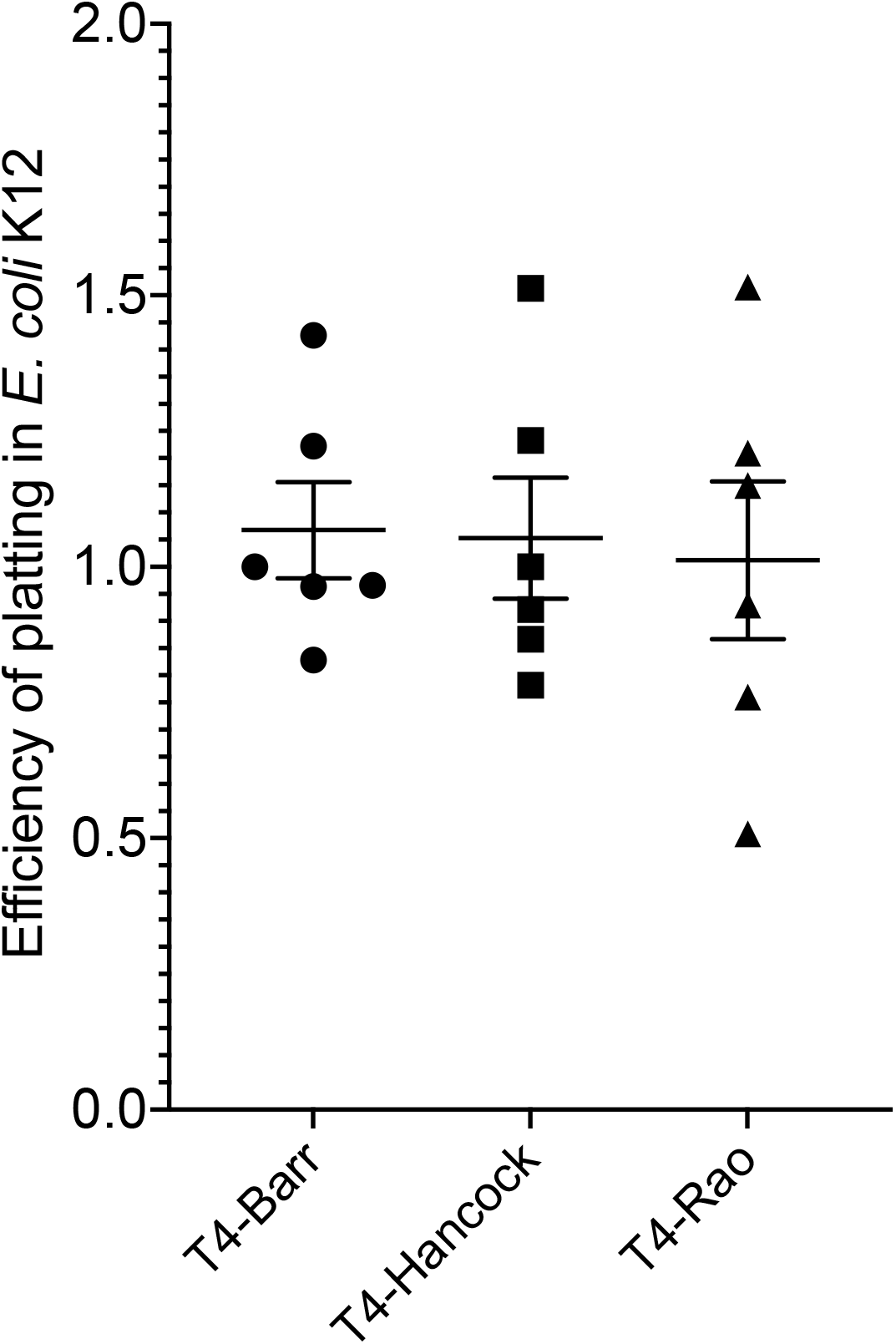
Efficiency of platting (EOP) for T4 sublines on *E. coli* K12. Each experiment was performed in duplicate and repeated three times on separate occasions. The values in the graph represent mean of six replicates and error bars represent standard error of the mean (p=0.91).

Next, we compared the replication cycle variables of latency period and burst size using one-step growth curves on the same pair of hosts. Our results revealed that all T4 sublines had similar growth features, characterised by a latent period of approximately 20 min, followed by a maturation period of ~10 min, reaching their plateau within ~30 min (Figure 4) (Ellis and Delbruck 1939). Similarly, the average burst size of all the T4 sublines were comparable within the two bacterial hosts. However, the burst size in *E. coli* B was calculated to be approximately 190 phage particles per infected bacterial cell, which was significantly higher than the average burst size of 109 phage particles per infected bacterial cell in *E. coli* K12.

**Figure 4:**
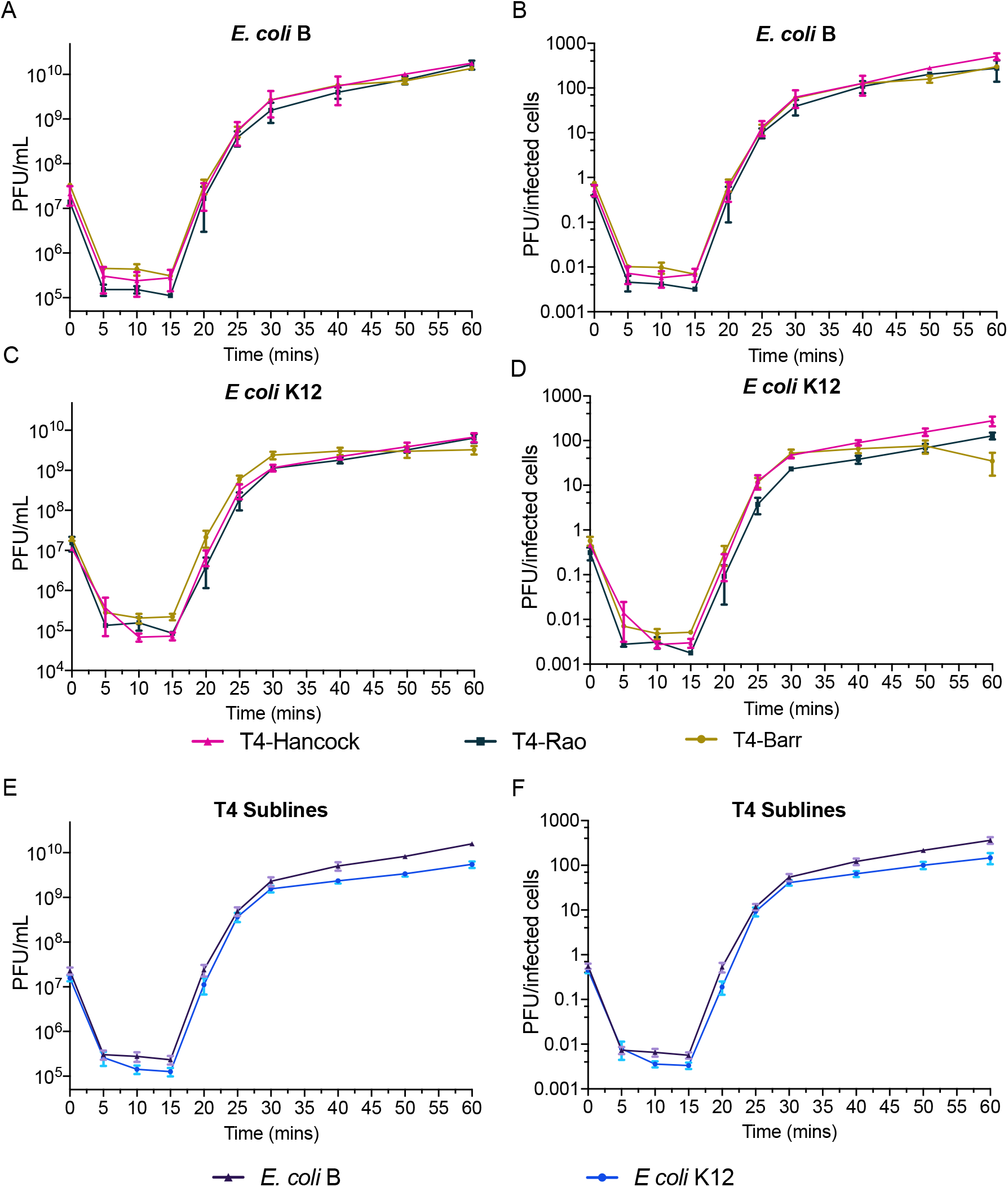
One step growth curve of T4 sublines in *E. coli* B (A and B) and *E. coli* K12 (C and D). The y-axis labelled PFU/infected cells (B and D) were calculated by dividing the plaque forming units in each time point by colony forming units of initial inoculum. Panels E and F represent average of all three T4 sublines in *E. coli* B and *E. coli* K12. All experiments were repeated in triplicate on three separate occasions. Error bars represent the standard error of mean.

Furthermore, to examine the rate of bacterial killing by phages, we performed growth kill assays. The three sublines of T4 were mixed at a multiplicity of infection (MOI) of 0.01 with actively growing (OD_600_ ~ 0.2) *E. coli* B and K12 and growth was measured by optical density (OD_600_) at 5 min intervals for 16 hours. All three sublines of T4 were able to suppress the growth of *E. coli* B and *E. coli* K12 within one hour, with a sharp reduction in OD observed in *E. coli* K12 with all three T4 sublines (Figure 5). The T4 sublines showed different growth patterns between the two different strains of *E. coli*, but there were no significant differences between the growth patterns between the T4 sublines.

**Figure 5:**
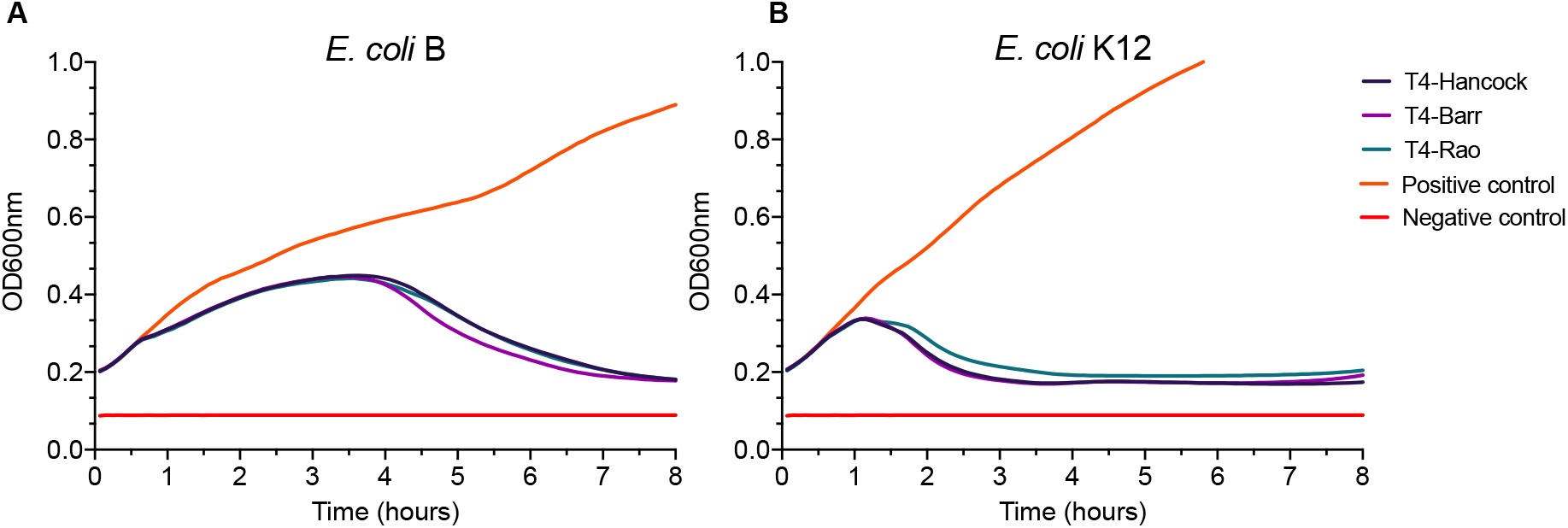
Growth kill curve of T4 sublines on *E. coli* B (A) and *E. coli* K12 (B) showing lytic activity of T4 phages for 8 hours. The optical density OD600nm was collected at 5 minutes interval and each point in the graph is the mean of thirty-six replicates (12 technical replicates in three biological repeats on separate occasions) (complete data is in supplementary figure 2).

Finally, we examined the host range of the T4-sublines on a selection of laboratory and pathogenic strains of *E. coli.* We used three standard laboratory strains and eight pathogenic strains for this assay. On these spot plate assays, we observed that T4 phages had lytic activity against six out of 11 strains, but there was no difference in host range between the T4-sublines (Table 4).

**Table 4.**
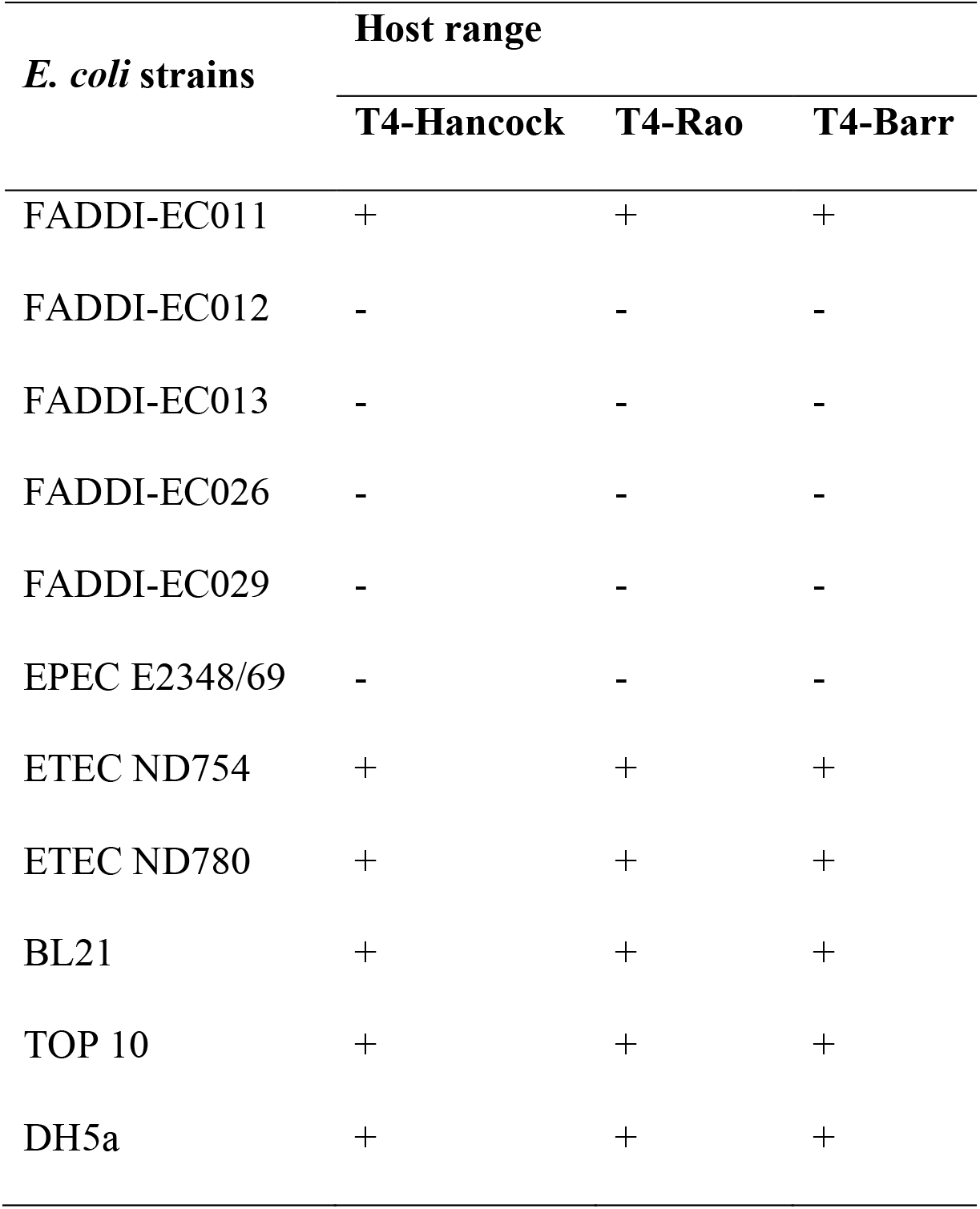
Host range of T4-sublines on laboratory and clinical strains of E. coli (+ = lysis on spot assay; − = no lysis on spot assay)

| *E. coli* strains | Host range |  |  |
| --- | --- | --- | --- |
|  | T4-Hancock | T4-Rao | T4-Barr |
| FADDI-EC011 | + | + | + |
| FADDI-EC012 | - | - | - |
| FADDI-EC013 | - | - | - |
| FADDI-EC026 | - | - | - |
| FADDI-EC029 | - | - | - |
| EPEC E2348/69 | - | - | - |
| ETEC ND754 | + | + | + |
| ETEC ND780 | + | + | + |
| BL21 | + | + | + |
| TOP 10 | + | + | + |
| DH5a | + | + | + |

## Discussion

T-series phages have been used as model systems to understand the fundamental molecular biology of life since the 1940s (Benzer 1955, Crick et al 1961, Jacob and Monod 1961, Luria and Delbruck 1943, Rath et al 2015, Sanger et al 1977, Smith et al 2003). Scientists throughout the world have been investigating the genetics and physiology of these bacteriophages, with many of the subsequent discoveries laying the foundation for the entire phage field. Given that these model phages have been distributed, maintained, and propagated across many different laboratories in a multitude of different ways, there exist the possibility for phenotypic and genotypic differences to accumulate between strains from different laboratories, which could ultimately compromise the comparability of these important model organisms.

To address this, we investigated the genetic and phenotypic changes between a 48-year old T-series phage stock and contemporary laboratory strains. Our analysis of three T4 sublines revealed minor genetic differences and no detectable variation in growth characteristic in their usual hosts *E. coli* B and *E. coli* K12. We did, however observe substantial differences between nucleotide sequence of NCBI RefSeq (Nucleotide accession no. AF158101) and our three T4 sublines, highlighting the need for an updated reference genome for T4 phage.

The complete genome of T4 was initially established through sequencing small fragments that were obtained following cloning and direct PCR (Miller et al 2003). According to the Genbank record, the first information on T4’s genome sequence was recorded in 1981 and its latest update was listed in 2003 (AF158101.6). A relatively high genetic variation between the Genbank RefSeq and T4 sublines of this study indicates that there may be artefacts in the original Genbank reference sequence. We therefore propose T4-Hancock as an updated reference genome for the field, and we used this genome as the reference in our analysis. The complete genome of T4-Hancock was 5 base-pair longer than the T4 Refseq. The reason for this difference in the genome length may be related to the fact that the genome of T4 is terminally redundant and circularly permuted, potentially altering the total length of chromosome between generations (Streisinger et al 1967).

We further examined the genetic divergence between historical and two contemporary stocks of T4 (T4-Barr and T4-Rao). Our analysis showed the insertion of 13 bp in T4-Barr and 14 bp in T4-Rao when compared with T4-Hancock. This finding is in line with total chromosome length of each subline that we obtained in our study. The average nucleotide identity percentage, although the difference was very small, showed that the two contemporary sublines; T4-Barr and T4-Rao shared a relatively higher percentage similarity to each other than they shared with T4-Hancock. Interestingly, both contemporary sublines also shared the nucleotide variations, with T4-Rao having five additional variations. The majority of nucleotide variations were in essential genes including genes that encode long tail fibers (gp34 and gp37), tail sheath (gp7) and enzymes associated with DNA metabolism. Although we could not track the history of our contemporary T4 phages, it is likely that these phages have gone through indefinite rounds of propagation, purification, and enumeration on various *E. coli* strains, the results of which may be associated with bottlenecks and mutational drift (Petrov et al 2010). Furthermore, the difference in mutation rate between T4-Rao and T4-Barr indicates that the mutational drift in T4 phages might have been driven by the different scale of work conducted between the laboratories, highlighting the possibility of genetic diversity of T4 sublines between laboratories across the world.

We, however, did not observe any noticeable differences in the growth characteristics between T4 sublines in our experimental conditions. Nevertheless, mutations in the essential structural genes, such as the long tail fibers, could potentially affect the phage’s adsorption to its host. T4 phage uses the C-terminal region of the distal subunit (gp37) of the long tail fibers to recognize host cellular receptors (Montag et al 1987, Montag et al 1990) and it has been shown that mutations in this region, which comprises about 140 amino acid residues in a 1026 residues-long polypeptide, could expand the host range of T4 and T4-like phages (Chen et al 2017, Tétart et al 1996). Our analysis showed that T4-Rao had a missense mutation in amino acid residue 417 of gp37, but we could not find any differences in host range between the T4-sublines on examination against eleven different laboratory and pathogenic strains of *E. coli.*

This study also examined the stability of T-series phages in prolonged storage. All the T-even series lysates had active phages present, which could be propagated in laboratory after 48 years in storage. However, we were not able to recover any viable phages from the lysate of T-odd series (T3 and T7). Although we do not have complete information on the long-term storage conditions of these phages over the last ~50 years, our results suggest that T-even series are more stable than T-odd series on prolonged storage. This finding is further supported by the close genetic relatedness of T-even phages, compared with the diversity seen across the T-odd phages (Abedon 2000).

In conclusion, our analyses suggest that individually maintained T4 sublines undergo continuous genetic drift that may cause micro-evolution in the model bacteriophage stocks. The genetic variation in T4 sublines did not show a difference in our phenotypic analysis, which is a favourable finding for the phage community, and the rationale proposed by Delbruck and co-workers on the use of model bacteriophages to avoid incomparability of results between laboratories proved to be still valid (Anderson 1992). However, the magnitude of genetic changes may vary between laboratories, which highlights a need for a larger-scale comparative study of model bacteriophages sublines.

## Acknowledgements

Authors would like to thank Prof. Peter Reeves, Michael Liu and Gordon Stevenson, from The University of Sydney, for the provision, shipping and long-term storage of the Hancock phage collection, Prof. Venigalla B. Rao, from The Catholic University of America, for providing a laboratory stock of T4 phage, Prof. Jian Li and Dr. Michael McDonald, Monash University and Roy M. Robins-Browne. University of Melbourne for providing *E. coli* strains and Dr. Fernando L Gordillo Altamirano, School of Biological Sciences, Monash University for editing the draft.

This work, including the efforts of Jeremy J. Barr, was funded by the Australian Research Council (ARC) Discovery Early Career Researcher Award (DECRA) (DE170100525), National Health and Medical Research Council (NHMRC: 1156588), and the Perpetual Trustees Australia award (2018HIG00007).

